# ADPriboDB v2.0: An Updated Database of ADP-ribosylated Proteins

**DOI:** 10.1101/2020.09.24.298851

**Authors:** Vinay Ayyappan, Ricky Wat, Calvin Barber, Christina A. Vivelo, Kathryn Gauch, Pat Visanpattanasin, Garth Cook, Christos Sazeides, Anthony K. L. Leung

**Affiliations:** Department of Biomedical Engineering, The G.W.C. Whiting School of Engineering, Johns Hopkins University, Baltimore, MD 21218, USA; Department of Biochemistry and Molecular Biology, Bloomberg School of Public Health, Johns Hopkins University, Baltimore, MD 21205, USA; Department of Biophysics, Krieger School of Arts and Sciences, Johns Hopkins University, Baltimore, MD 21218, USA; Department of Biology, Krieger School of Arts and Sciences, Johns Hopkins University, Baltimore, MD 21218, USA; Department of Molecular Biology and Genetics, School of Medicine, Johns Hopkins University, Baltimore, MD 21205, USA; Department of Oncology, School of Medicine, Johns Hopkins University, Baltimore, MD 21205, USA

## Abstract

ADP-ribosylation is a protein modification responsible for biological processes such as DNA repair, RNA regulation, cell cycle, and biomolecular condensate formation. Dysregulation of ADP-ribosylation is implicated in cancer, neurodegeneration, and viral infection. We developed ADPriboDB (adpribodb.leunglab.org) to facilitate studies in uncovering insights into the mechanisms and biological significance of ADP-ribosylation. ADPriboDB 2.0 serves as a one-stop repository comprising 48,346 entries and 9,097 ADP-ribosylated proteins, of which 6,708 were newly identified since the original database release. In this updated version, we provide information regarding the sites of ADP-ribosylation in 32,946 entries. The wealth of information allows us to interrogate existing databases or newly available data. For example, we found that ADP-ribosylated substrates are significantly associated with the recently identified human protein interaction networks associated with SARS-CoV-2, which encodes a conserved protein domain called macrodomain that binds and removes ADP-ribosylation. In addition, we create a new interactive tool to visualize the local context of ADP-ribosylation, such as structural and functional features as well as other post-translational modifications (e.g., phosphorylation, methylation and ubiquitination). This information provides opportunities to explore the biology of ADP-ribosylation and generate new hypotheses for experimental testing.

## Introduction

ADP-ribosylation is a reversible post-translational modification defined by the addition of one [mono(ADP-ribosyl)ation] or multiple [poly(ADP-ribosyl)ation] ADP-ribose moieties onto amino acid side chains with nucleophilic oxygen, nitrogen, or sulfur (1,2). ADP ribose groups can be transferred by enzymatically active members of the family of 17 ADP-ribosyltransferases (commonly known as PARPs) as well as other enzymes, such as bacterial toxins and sirtuins (3–5). ADP-ribosylation has been implicated in fundamental biological processes (e.g., DNA damage repair, gene regulation, and cell signaling) (6–8) as well as disease-related processes (e.g., microbial pathogenesis, carcinogenesis, and inflammation) (9–11). While ADP-ribosylation was discovered in 1963 (12), studies of ADP-ribosylation have recently begun to benefit from the rapid development in proteomics technologies that identify ADP-ribosylated sites and characterize the substrate specificity of PARPs and other enzymes associated with ADP-ribosylation (13). In light of these developments, we created ADPriboDB in 2015 (14) to provide a one-stop informatics portal about ADP-ribosylated substrates across the proteomes from different species, similar to databases like dbPTM (15, 16) and PhosphoSite (17) that provide information about other post-translational modifications such as ubiquitination and phosphorylation. In addition, ADPriboDB provides data entries pertaining to PARP inhibitor effects on the ADP-ribosylated proteome, aiming to bridge basic scientific knowledge about ADP ribosylation with information that physicians can use to garner insights on clinical benefits and side effects.

ADPriboDB 2.0 includes an expanded library of data, new addition of site information, and an improved user interface (**Table 1**). ADPriboDB 2.0 also integrates other post-translational modification sites and features a web interface that allows users to visualize protein domains and other features at and proximal to the modification sites.

**Table 1:**
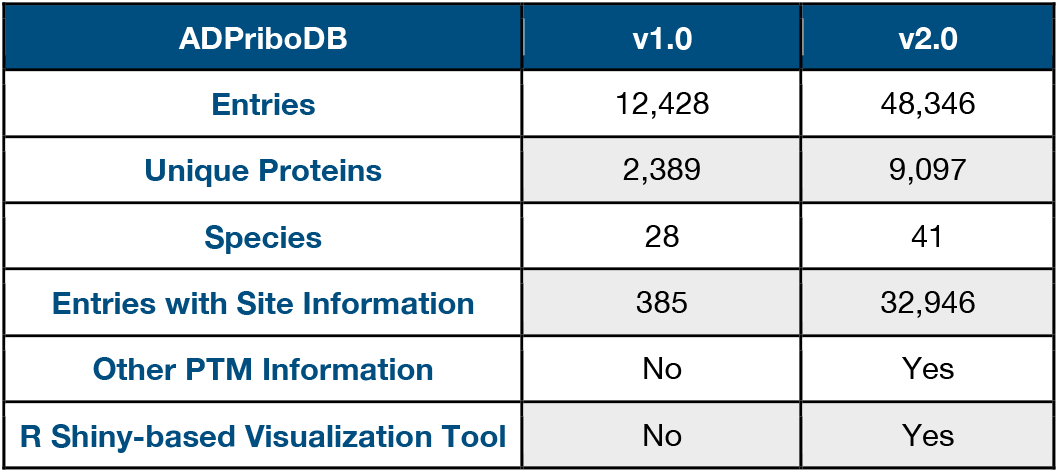
Data summary of ADPriboDB version

## New Features

### Expanded Library of Data

As in ADPriboDB, published papers were curated from PubMed, accessed via the following search terms: ‘PARylation’ or ‘PARsylation’ or ‘poly(ADP-ribosyl)ation’ or ‘poly-adp-ribosylation’ or ‘poly(ADP-ribosylation)’ or ‘PARylated’ or ‘PARsylated’ or ‘poly(ADP)ribosylation’ or ‘MARylation’ or ‘MARsylation’ or ‘mono(ADP-ribosyl)ation’ or ‘mono-adp-ribosylation’ or ‘mono(ADP-ribosylation)’ or ‘MARylated’ or ‘MARsylated’ or ‘mono(ADP)ribosylated.’ Search results were restricted between the dates of July 2015 and May 2019, where the initial release covered between January 1975 and June 2015 (14). ADPriboDB 2.0 now also included the terms ‘ADPr,’ ‘ADP-ribose,’ and ‘ADPribosylation’ for literature search between January 1975 and May 2019. Information regarding protein identifiers as well as experimental and modification details were collected and assessed for inclusion in ADPriboDB by two independent curators.

Since 2010, research regarding ADP-ribosylation has accelerated, and particularly the identification of ADP-ribosylated substrates has risen (**Figure 1A**). In light of these trends, the latest iteration of ADPriboDB comprises 48,346 entries, including 9,097 unique proteins across 610 papers, more than tripling the size of the initial release of the database. The database now includes data from 41 species, up from 28 species. ADPriboDB 2.0 has more than doubled the proportion of its entries bearing information regarding the enzyme responsible for the ADP-ribosylation. Reflecting a greater availability of technology (e.g. analog-sensitive PARP approaches), 65% of database entries include the information of enzyme-substrate specificity (**Table 2**). To enhance the speed with which queried database pages are loaded, we have used the open-source PHP framework Xataface to compress and save all requested pages to a cache. These caches are generated for each individual user. This feature allows the display of the frequently visited pages, thereby speeding-up the process of information loading and display.

**Figure 1.**
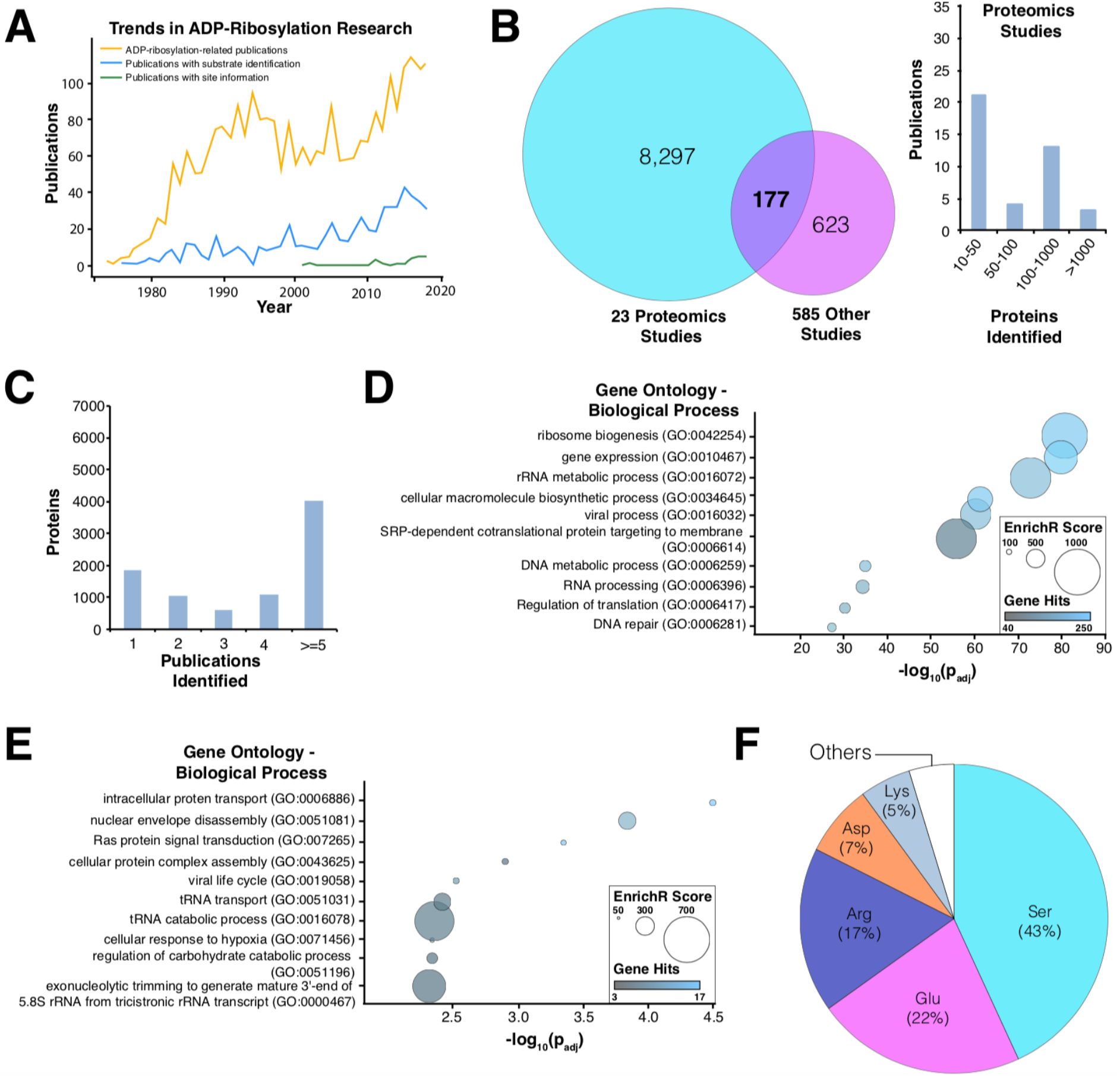
Analyses of ADPriboDB v2.0. (**A**) Trends in ADP-ribosylation research shows a progressive increase in the number of publications about ADP-ribosylation biology, substrates and sites. (**B**) Left: Venn diagram shows the overlap between proteins identified in proteomics-based studies (≥10 proteins) and those identified in other publications. Right: Histogram describes the number of proteins identified by these proteomics studies. (**C**) Distribution of the number of publications identifying a given ADP-ribosylated substrate. (**D**) Proteins identified in at least two publications were subjected to gene ontology analysis via EnrichR. Bubble chart shows the enrichment of selected pathways analyzed with REVIGO (26); the full enrichment analysis is available in Supplementary Data S1. (**E**) Enrichment analysis was also performed on ADP-ribosylated proteins within the SARS-CoV-2 interactome. Bubble chart shows the enrichment of selected pathways analyzed with REVIGO; the full enrichment analysis is available in Supplementary Data S2. (**F**) Distribution of ADP-ribosylated residues.

**Table 2:** Number of unique proteins modified by each PARP enzyme

| PARP Enzyme | Modified Proteins |
| --- | --- |
| PARP1 | 1,524 |
| PARP10 | 1,080 |
| PARP2 | 462 |
| PARP3 | 394 |
| PARP11 | 295 |
| PARP14 | 222 |
| Other PARPs | 58 |

In ADPriboDB 2.0, a growing number of entries were derived from proteomics studies, which account for ~8,000 unique proteins (**Figure 1B**). ~75% of ADP-ribosylated proteins were identified in at least two publications (**Figure 1C**). Analysis of these independently verified ADP-ribosylated substrates via EnrichR (18) revealed a significant enrichment of gene sets associated with diverse biological processes, including DNA metabolism, RNA processing, protein targeting, and viral-related processes (**Figure 1D**; **Supplementary Data S1**).

This expanded database therefore provides greater flexibility and depth to analyses of ADP-ribosylation, its function, and its clinical implications. As an example, we used ADPriboDB to explore the biology of the severe acute respiratory syndrome coronavirus 2 (SARS-CoV-2). SARS-CoV-2 encodes a protein domain (called macrodomain) that is conserved in all coronaviruses (19, 20), and this macrodomain binds and hydrolyzes ADP-ribose from substrates (21, 22), as in other viral macrodomains. Given the conserved nature of macrodomain and the critical importance of the enzymatic activity of macrodomain in the virulence of other coronaviruses (19, 20), ADP-ribosylation is likely highly regulated in SARS-CoV-2 infection. Consistent with this hypothesis, human protein interactors of SARS-CoV-2 viral proteins (23) are statistically enriched with ADP-ribosylated substrates (p=0.0096, Chi-square test with continuity correction) (**Supplementary Data S2**). No statistical significance was observed between SARS-CoV-2 protein interactors with the methylated proteomes (p=0.1977, Chi-square test with continuity correction). Notably, this statistical enrichment with ADP-ribosylated substrates was present even though the SARS-CoV-2 protein interaction map did not include the association with nonstructural protein 3 (nsP3), which possesses the macrodomain (23). These ADP-ribosylated substrates associated with SARS-CoV-2 proteins were enriched with gene ontologies, including transfer RNA and ribosomal RNA regulation (**Figure 1E, Supplementary Data S2**).

### Increased Availability of ADP-ribosylation Site Information

For the last six years, evolving proteomics techniques have allowed the identification of sites of ADP-ribosylation for functional analyses (13, 24). During curation of database entries, information regarding modified sites and/or peptide sequences identified by mass spectrometry were included when available. Sequence information was then aligned to protein sequences deposited in the UniProt database (25). Currently, ~67% of ADPriboDB 2.0 entries includes information regarding ADP-ribosylation sites, with a total of 14,839 unique sites on a range of amino acids (**Figure 1F**). For each database entry, users can view the site and sequence information provided by the publication. Users can also find out whether the same sites were identified in other publications or whether there are other ADP-ribosylation sites from the same proteins.

### Enhanced User Interface Features

The new site information enables users to identify potential crosstalks between ADP-ribosylation and other post-translational modifications. ADPriboDB 2.0 currently features sites of phosphorylation, methylation, and ubiquitination, sourced from dbPTM (15, 16). To facilitate correlation analyses, ADPriboDB 2.0 includes a new Shiny-based tool to visualize the local structural and functional context of ADP-ribosylation sites, such as other post-translational modification sites, protein domains, as well as regions of protein-protein interaction. Vertical black lines are drawn to denote the location of ADP-ribosylation sites. For example, a high frequency of Axin1 ADP-ribosylation was found to localize around the motif that binds tankyrases—i.e., PARP5a and PARP5b (**Figure 2A**).

**Figure 2:**
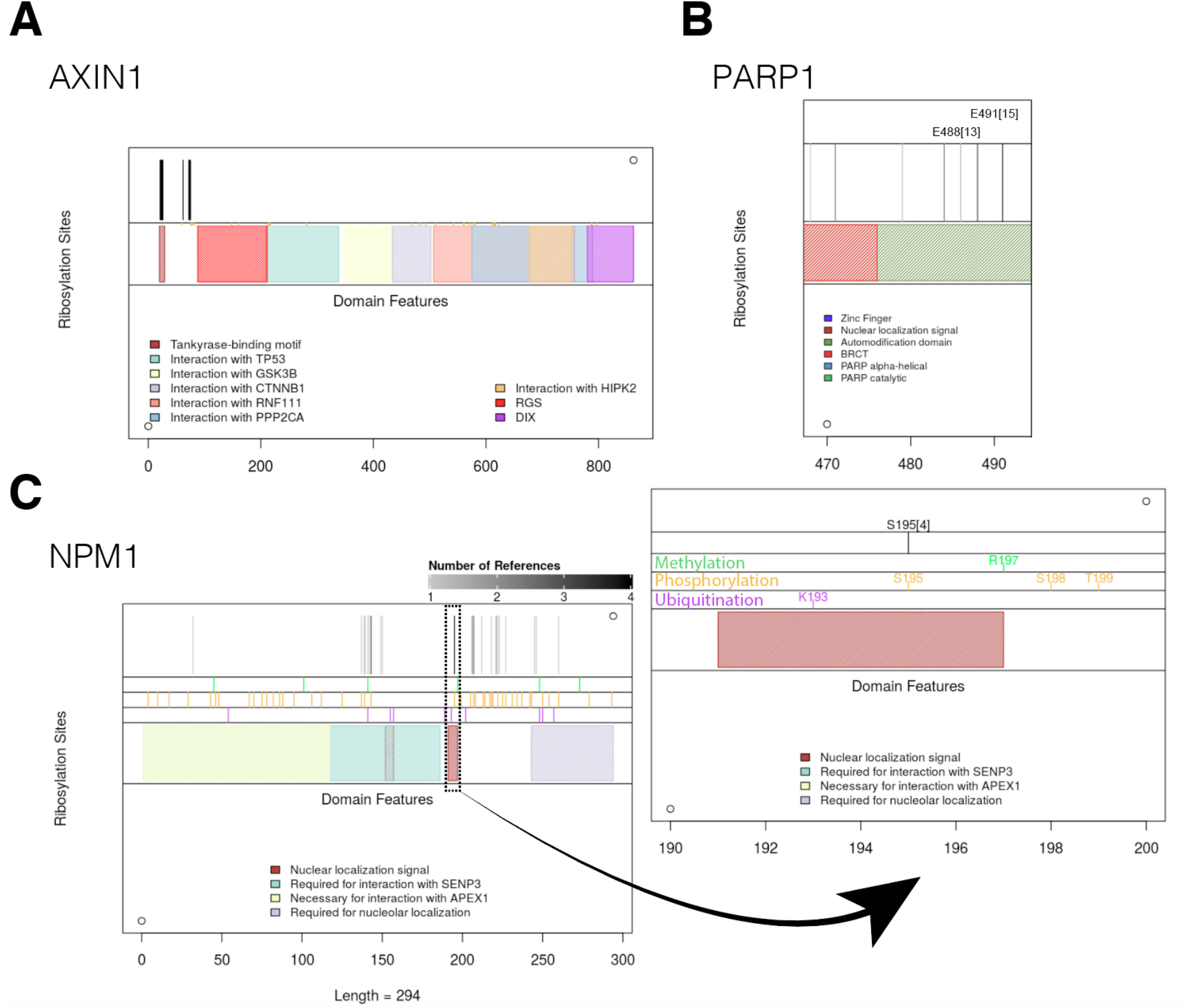
New Graphical Interface. Representative images shown for (**A**) AXIN1 (all sites have one reference) (**B**) PARP1, and (**C**) NPM1.

To help identify ADP-ribosylation sites of high confidence, the number of publications that identified a given modified site is now provided. So far, ~7% of sites (1,018 unique sites) listed in the database have been identified by at least two publications (**Supplementary Figure S1**). The varying grey level of the vertical black lines corresponds to the number of publications identifying that particular site in a given protein. Users can then gauge the “confidence” of a particular ADP-ribosylation site for mutagenesis studies. For example, E488 and E491 within the automodification domain of PARP1 were identified in more than 10 independent studies (**Supplementary Data S3, Figure 2B**).

Besides, users can toggle zoom settings to visualize crowded regions at a single amino acid level (**Figure 2C**). For example, S195 in NPM1 within the nuclear localization signal were identified 4 times, and this site can also be phosphorylated (**Figure 2C**). Taken together, all these additional features enable users to prioritize which sites for further mechanistic analyses.

### Conclusions and Future Development

ADPriboDB v2.0 was updated in response to the growing volume of data regarding novel substrates and sites of ADP-ribosylation. We anticipate the wealth of site information will stimulate new mechanistic studies, especially with the advent of new chemical approaches to generate model ADP-ribosylated peptides and methods to label ADP-ribose (27, 28). Given that the database now includes a considerable amount of site and sequence information, we hope the database will encourage the development of novel algorithms for motif finding, site prediction, and correlation analyses with various post-translational modifications (29, 30).

With the development of chemical inhibitors and genetic ablation technologies (such as CRISPR) that can target specific PARPs, we anticipate future generations of ADPriboDB will include more information regarding substrate-enzyme specificity. As ADP-ribosylhydrolases also exhibit amino acid specificity (31), future entries may include information of enzymes that add *and* remove ADP-ribosylation from specific sites. Emergent studies indicate that cellular processes are not only regulated by the balance between the synthesis and removal of ADP-ribosylation, but also by maintaining the appropriate forms of ADP-ribosylation (32). Although mono(ADP-ribosyl)ation and poly(ADP-ribosyl)ation cannot be distinguished by the current proteomics methods (13), future entries will be focused on annotating this critical chain length information, especially when appropriate technologies are mature (28, 33).

## Data Availability

ADPriboDB is free and publicly available at http://adpribodb.leunglab.org/

## Supplementary Data

**Supplementary Data S1:** Enrichment analysis of proteins identified to be ADP-ribosylated in at least two publications based “Gene Ontology: Biological Process” pathways.

**Supplementary Data S2:** Enrichment analysis of ADP-ribosylated that interacts with SARS-CoV-2 proteins based on “Gene Ontology: Biological Process” pathways.

**Supplementary Data S3:** Domains in which ADP-ribosylated residues are found, among the sites identified as ADP-ribosylated by at least two publications.

Supplementary Data are available at NAR online.

## Acknowledgements

We thank the Leung lab members for critical comments on the manuscript and the database for improving its usability.

---

**Supplementary Figure S1.**
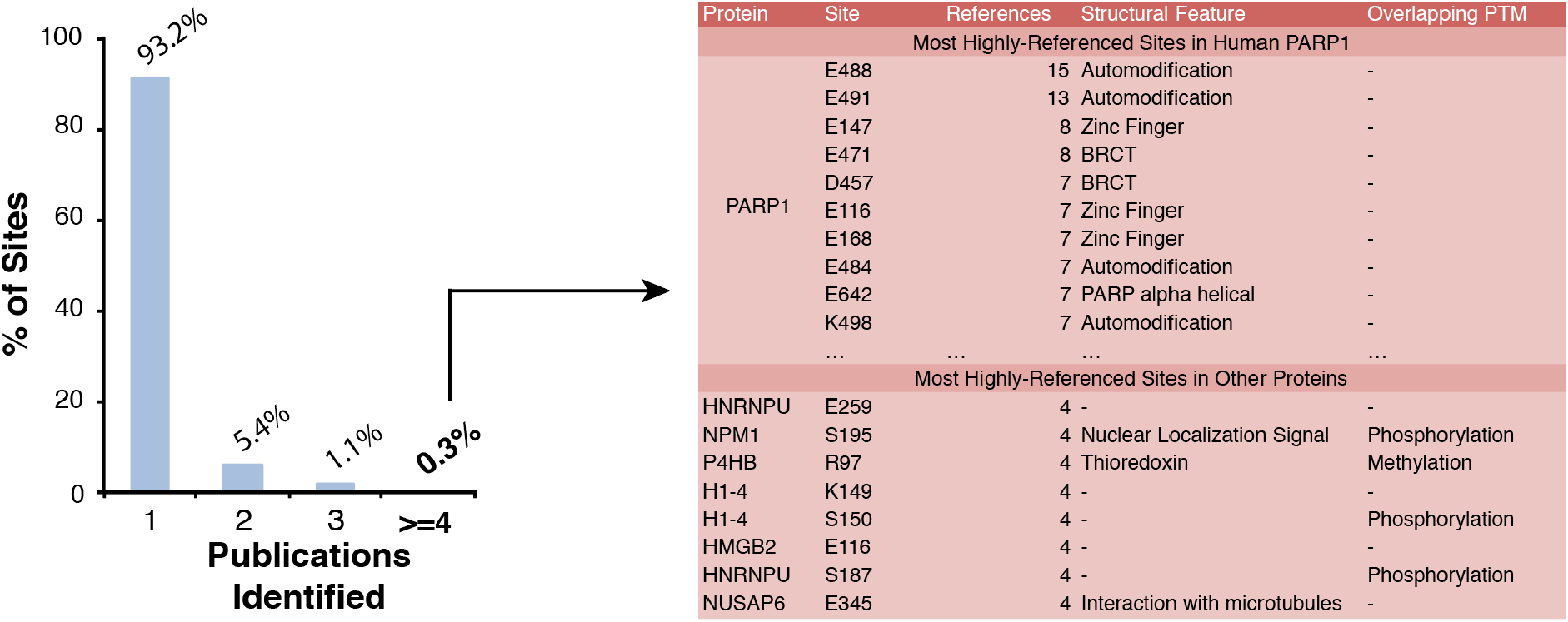
Analysis of ADP-ribosylation sites. **(A)** Left: Boxplot indicate the percentage of ADP-ribosylation sites identified by the number of publications. Right: Shown are sites identified in at least four publications. **(B)** Amino acid distribution among sites identified by >=2 and 1 publications, respectively.

## References

1. Gupte, R., Liu, Z. and Kraus, W.L. (2017) PARPs and ADP-ribosylation: recent advances linking molecular functions to biological outcomes. Genes Dev., 31, 101–126.

2. Palazzo, L., Mikoč, A. and Ahel, I. (2017) ADP-ribosylation: new facets of an ancient modification. FEBS J., 284, 2932–2946.

3. Houtkooper, R.H., Pirinen, E. and Auwerx, J. (2012) Sirtuins as regulators of metabolism and healthspan. Nat. Rev. Mol. Cell Biol., 13, 225–238.

4. Hottiger, M.O., Hassa, P.O., Lüscher, B., Schüler, H. and Koch-Nolte, F. (2010) Toward a unified nomenclature for mammalian ADP-ribosyltransferases. Trends Biochem. Sci., 35, 208–219.

5. Di Girolamo, M., Dani, N., Stilla, A. and Corda, D. (2005) Physiological relevance of the endogenous mono(ADP-ribosyl)ation of cellular proteins. FEBS J., 272, 4565–4575.

6. Hottiger, M.O. (2015) Nuclear ADP-Ribosylation and Its Role in Chromatin Plasticity, Cell Differentiation, and Epigenetics. Annu. Rev. Biochem., 84, 227–263.

7. Ray Chaudhuri, A. and Nussenzweig, A. (2017) The multifaceted roles of PARP1 in DNA repair and chromatin remodelling. Nat. Rev. Mol. Cell Biol., 18, 610–621.

8. Kim, D.-S., Challa, S., Jones, A. and Kraus, W.L. (2020) PARPs and ADP-ribosylation in RNA biology: from RNA expression and processing to protein translation and proteostasis. Genes Dev., 34, 302–320.

9. Simon, N.C., Aktories, K. and Barbieri, J.T. (2014) Novel bacterial ADP-ribosylating toxins: structure and function. Nat. Rev. Microbiol., 12, 599–611.

10. Hou, W.-H., Chen, S.-H. and Yu, X. (2019) Poly-ADP ribosylation in DNA damage response and cancer therapy. Mutat. Res., 780, 82–91.

11. Fehr, A.R., Singh, S.A., Kerr, C.M., Mukai, S., Higashi, H. and Aikawa, M. (2020) The impact of PARPs and ADP-ribosylation on inflammation and host-pathogen interactions. Genes Dev., 34, 341–359.

12. Chambon, P., Weill, J.D. and Mandel, P. (1963) Nicotinamide mononucleotide activation of new DNA-dependent polyadenylic acid synthesizing nuclear enzyme. Biochem. Biophys. Res. Commun., 11, 39–43.

13. Daniels, C.M., Ong, S.-E. and Leung, A.K.L. (2015) The promise of proteomics for the study of ADP-ribosylation. Mol. Cell, 58, 911–924.

14. Vivelo, C.A., Wat, R., Agrawal, C., Tee, H.Y. and Leung, A.K.L. (2016) ADPriboDB: The database of ADP-ribosylated proteins. Nucleic Acids Res., 45, D204–D209.

15. Huang, K.-Y., Su, M.-G., Kao, H.-J., Hsieh, Y.-C., Jhong, J.-H., Cheng, K.-H., Huang, H.-D. and Lee, T.-Y. (2016) dbPTM 2016: 10-year anniversary of a resource for post-translational modification of proteins. 44, D435–46.

16. Huang, K.-Y., Lee, T.-Y., Kao, H.-J., Ma, C.-T., Lee, C.-C., Lin, T.-H., Chang, W.-C. and Huang, H.-D. (2019) dbPTM in 2019: exploring disease association and cross-talk of post-translational modifications. Nucleic Acids Res., 47, D298–D308.

17. Hornbeck, P.V., Zhang, B., Murray, B., Kornhauser, J.M., Latham, V. and Skrzypek, E. (2015) PhosphoSitePlus, 2014: mutations, PTMs and recalibrations. 43, D512–20.

18. Kuleshov, M.V., Jones, M.R., Rouillard, A.D., Fernandez, N.F., Duan, Q., Wang, Z., Koplev, S., Jenkins, S.L., Jagodnik, K.M., Lachmann, A., et al. (2016) Enrichr: a comprehensive gene set enrichment analysis web server 2016 update. Nucleic Acids Research, 44, W90–W97.

19. Leung, A.K.L., McPherson, R.L. and Griffin, D.E. (2018) Macrodomain ADP-ribosylhydrolase and the pathogenesis of infectious diseases. PLoS Pathog., 14, e1006864.

20. Alhammad, Y.M.O. and Fehr, A.R. (2020) The Viral Macrodomain Counters Host Antiviral ADP-Ribosylation. Viruses, 12.

21. Frick, D.N., Virdi, R.S., Vuksanovic, N., Dahal, N. and Silvaggi, N.R. (2020) Molecular Basis for ADP-Ribose Binding to the Mac1 Domain of SARS-CoV-2 nsp3. Biochemistry, 59, 2608–2615.

22. Alhammad, Y.M.O., Kashipathy, M.M., Roy, A., Gagné, J.-P., Nonfoux, L., McDonald, P., Gao, P., Battaile, K.P., Johnson, D.K., Poirier, G.G., et al. The SARS-CoV-2 conserved macrodomain is a highly efficient ADP-ribosylhydrolase. 10.1101/2020.05.11.089375.

23. Gordon, D.E., Jang, G.M., Bouhaddou, M., Xu, J., Obernier, K., White, K.M., O’Meara, M.J., Rezelj, V.V., Guo, J.Z., Swaney, D.L., et al. (2020) A SARS-CoV-2 protein interaction map reveals targets for drug repurposing. Nature, 583, 459–468.

24. Zhen, Y. and Yu, Y. (2018) Proteomic Analysis of the Downstream Signaling Network of PARP1. Biochemistry, 57, 429–440.

25. UniProt Consortium (2019) UniProt: a worldwide hub of protein knowledge. Nucleic Acids Res., 47, D506–D515.

26. Supek, F., Bošnjak, M., Škunca, N. and Šmuc, T. (2011) REVIGO summarizes and visualizes long lists of gene ontology terms. PLoS One, 6, e21800.

27. van der Heden van Noort, G.J. (2020) Chemical Tools to Study Protein ADP-Ribosylation. ACS Omega, 10.1021/acsomega.9b03591.

28. Ando, Y., Elkayam, E., McPherson, R.L., Dasovich, M., Cheng, S.-J., Voorneveld, J., Filippov, D.V., Ong, S.-E., Joshua-Tor, L. and Leung, A.K.L. (2019) ELTA: Enzymatic Labeling of Terminal ADP-Ribose. Mol. Cell, 73, 845–856.e5.

29. Li, G.X.H., Vogel, C. and Choi, H. (2018) PTMscape: an open source tool to predict generic post-translational modifications and map modification crosstalk in protein domains and biological processes. Mol Omics, 14, 197–209.

30. Chang, C.-C., Tung, C.-H., Chen, C.-W., Tu, C.-H. and Chu, Y.-W. (2018) SUMOgo: Prediction of sumoylation sites on lysines by motif screening models and the effects of various post-translational modifications. Sci. Rep., 8, 15512.

31. Rack, J.G.M., Palazzo, L. and Ahel, I. (2020) (ADP-ribosyl)hydrolases: structure, function, and biology. Genes Dev., 34, 263–284.

32. Suskiewicz, M.J., Zobel, F., Ogden, T.E.H., Fontana, P., Ariza, A., Yang, J.-C., Zhu, K., Bracken, L., Hawthorne, W.J., Ahel, D., et al. (2020) HPF1 completes the PARP active site for DNA damage-induced ADP-ribosylation. Nature, 579, 598–602.

33. Martello, R., Mangerich, A., Sass, S., Dedon, P.C. and Bürkle, A. (2013) Quantification of cellular poly(ADP-ribosyl)ation by stable isotope dilution mass spectrometry reveals tissue-and drug-dependent stress response dynamics. ACS Chem. Biol., 8, 1567–1575.

